# Solution structure, dynamics and tetrahedral assembly of Anti-TRAP, a homo-trimeric triskelion-shaped regulator of tryptophan biosynthesis in *Bacillus subtilis*

**DOI:** 10.1101/2023.06.29.547145

**Authors:** Craig McElroy, Elihu Ihms, Deepak Kumar Yadav, Melody Holmquist, Vibhuti Wadwha, Vicki Wysocki, Paul Gollnick, Mark Foster

## Abstract

Cellular production of tryptophan is metabolically expensive and tightly regulated. The small *Bacillus subtilis* zinc binding Anti-TRAP protein (AT), which is the product of the *yczA/rtpA* gene, is upregulated in response to accumulating levels of uncharged tRNA^Trp^ through a T-box antitermination mechanism. AT binds to the undecameric ring-shaped protein TRAP (*trp* RNA Binding Attenuation Protein), thereby preventing it from binding to the *trp* leader RNA. This reverses the inhibitory effect of TRAP on transcription and translation of the *trp* operon. AT principally adopts two symmetric oligomeric states, a trimer (AT_3_) featuring a three-helix bundle, or a dodecamer (AT_12_) comprising a tetrahedral assembly of trimers, whereas only the trimeric form has been shown to bind and inhibit TRAP. We demonstrate the utility of native mass spectrometry (nMS) and small-angle x-ray scattering (SAXS), together with analytical ultracentrifugation (AUC) for monitoring the pH and concentration-dependent equilibrium between the trimeric and dodecameric structural forms of AT. In addition, we report the use of solution nuclear magnetic resonance (NMR) spectroscopy to determine the solution structure of AT_3_, while heteronuclear ^15^N relaxation measurements on both oligomeric forms of AT provide insights into the dynamic properties of binding-active AT_3_ and binding-inactive AT_12_, with implications for TRAP inhibition.

## Introduction

The *trp* RNA-binding attenuation protein (TRAP) senses intracellular levels of tryptophan in order to regulate transcription [1–4] and translation [5–10] of the genes required for tryptophan biosynthesis (Figure 1). Transcriptional and translational control by TRAP depends on the ability of tryptophan-bound TRAP to bind to an RNA site with multiple (G/U)AG repeats separated by two or three nonconserved spacer nucleotides [1]. The transcription of the six tryptophan biosynthetic genes clustered in the *trpECDFBA* operon [11] is controlled through an attenuation mechanism by which TRAP binding is thought to remodel competing secondary structural elements in the 5′ leader region of the nascent mRNA: in the absence of tryptophan, RNA binding by TRAP is impaired, and the leader folds into an anti-terminator hairpin that allows transcription of the entire operon. Increased cellular tryptophan binds to TRAP, activating it to bind to its RNA target which overlaps with and prevents formation of the anti-terminator hairpin; instead, a small downstream terminator hairpin forms and transcription of the structural genes is attenuated [2]. There is considerable evidence in support of this terminator/anti-terminator model, although TRAP nevertheless induces attenuation in the absence of an intact terminator [12,13].

**Figure 1.**
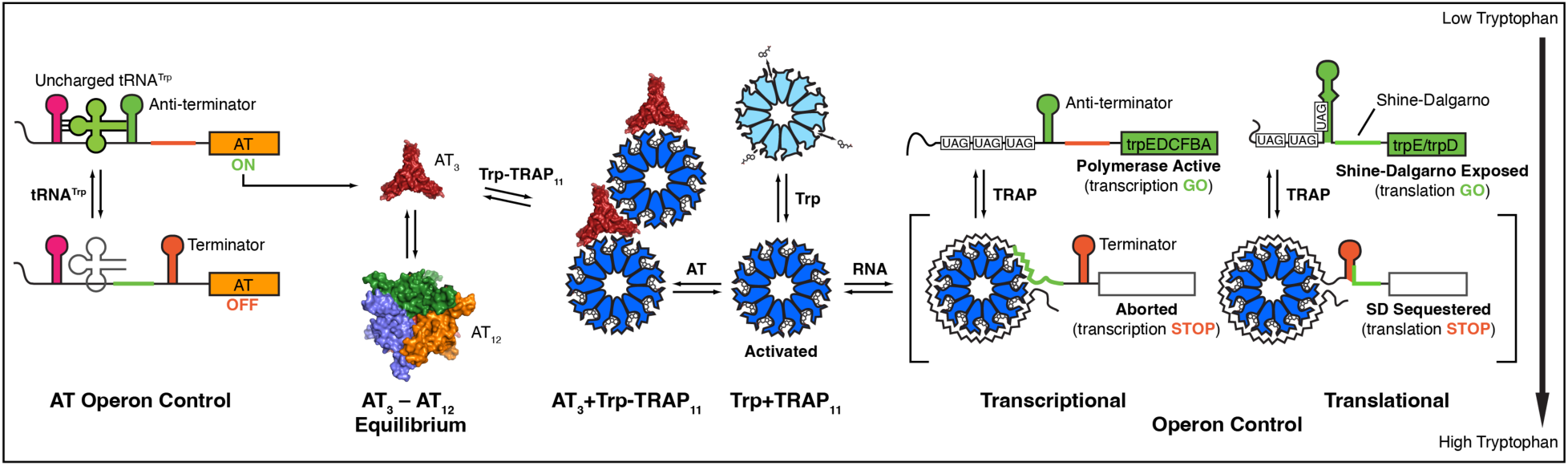
Transcription and translation of the *trp* operon in *B. subtilis* is regulated by feedback loops involving the functions of two proteins: the ring-shaped 11-mer TRAP, which senses free Trp, and Anti-TRAP (AT) which is overexpressed in response to accumulation of uncharged tRNA^Trp^. In the absence of AT, Trp-bound TRAP (blue) binds a series of UAG and GAG triplets in the 5’ leader of the *trp* operon, resulting in transcriptional and translational repression in part by formation of a terminator or sequestering the Shine-Dalgarno (SD) sequence. When expressed, trimeric AT (AT3) can bind to Trp-activated TRAP, preventing RNA binding and blocking its inhibition of Trp biosynthesis. AT exists in equilibrium between trimeric (red) and dodecameric states (blue, orange, green; the red trimer is behind the other three), while only trimeric AT binds and inhibits TRAP.

Translational control of *trpE* [9,14], and perhaps the entire *trp* operon, is also accomplished through TRAP binding-dependent RNA secondary structure [8]. When TRAP is inactive and cannot bind to the read-through transcript of the *trp* operon, the Shine-Dalgarno (SD) sequence of the *trpE* gene is unstructured and allows ribosome binding. When Trp-activated TRAP binds to the untranslated region of the transcript, the *trpE* SD sequence becomes sequestered in a hairpin that forms downstream of the TRAP binding site, thereby preventing ribosome binding and translation initiation [9]. TRAP binding sites are also present in the transcripts of the *trpG*, *trpP*, and *ycbK* genes, but in the translation initiation region, where TRAP binding directly impedes ribosome binding and thereby regulates translation [1,6,15,16].

In many bacteria, the extent of charging of tRNA^Trp^ is sensed as a regulatory signal in addition to the levels of free tryptophan [17,18]. In studies with a temperature-sensitive mutant of the tryptophanyl-tRNA synthetase in *B. subtilis*, the accumulation of uncharged tRNA^Trp^ leads to over-expression of the *trpECDFBA* operon [15]. The operon responsible for this effect was shown to be regulated via the T-box transcription anti-termination mechanism, where uncharged tRNA^Trp^ specifically pairs with the leader RNA causing anti-termination and transcriptional read-through [6]. Deletions within the operon identified *yczA/rtpA* as the responsible gene, and the protein product, anti-TRAP (AT), was shown to inhibit TRAP activity [6,19,20]. Expression of AT was also shown to be regulated at the level of translation by tRNA-dependent ribosome stalling [21,22].

Symmetry mismatch between the oligomeric states of AT and TRAP was evident from early chemical cross-linking, analytical ultracentrifugation, native mass spectrometry and NMR studies [19,23–26]. Diffracting crystals of *Bsu* AT revealed a dodecamer (AT_12_) arranged as a tetrahedral tetramer of trimers (AT_3_)_4_ (Figure 2) [27–29]. Each AT protomer forms a short helix (residues 5-8) at the N-terminus, β-strands from residues 9-11 and 34-36 form a short anti-parallel ý-sheet; a β-hairpin is formed by strands from residues 20-21 and 24-25, and an α-helix is present at the C-terminus from residues 38-50. Trimers are arranged via a three-helix bundle involving the C-terminal helix, with the zinc binding domains extended outward from the C_3_ helical axis. The AT dodecamer is formed through complementary inter-trimer ion pair interactions involving an N-terminal amine and a sidechain carboxylate from each trimer, as well as hydrophobic surface burial of the zinc-binding regions (Figure 2) [27–29]. Because the stabilizing ion pair requires a protonated N-terminus, which has a pKa near pH 7 [30], assembly of the dodecamer is disfavored above pH ∼8 [26]. The *B. licheniformis* variant of AT (which shares a 76% sequence identity with *B. subtilis* AT) crystallized at low pH adopts the same dodecameric structure as the *Bsu* protein, but at high pH the trimers assemble in an inverted orientation, with their N-termini projecting into solvent [29].

**Figure 2.**
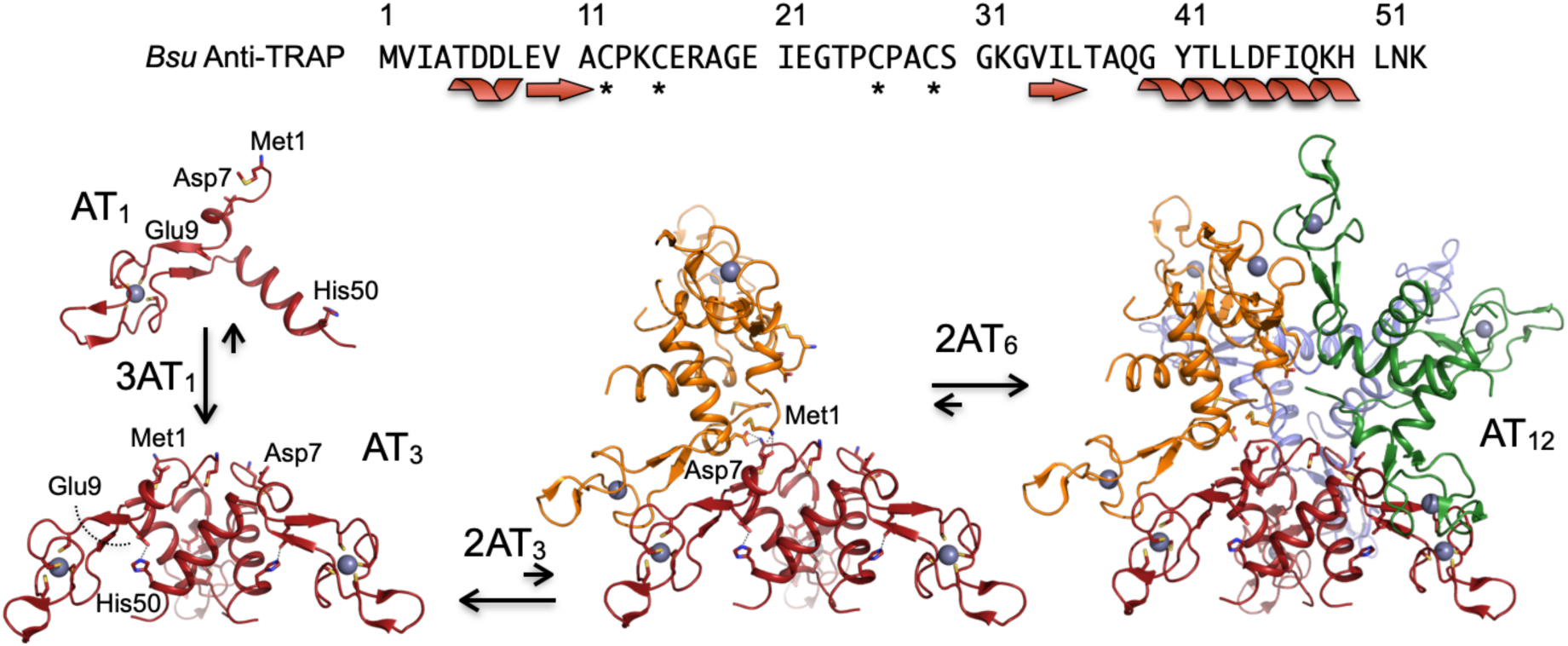
Oligomeric assembly of *Bsu* Anti-TRAP (AT). The 53-residue AT protomer (AT1) features a short N-terminal helix leading into a “zinc ribbon” motif and ending in a C-terminal helix. Assembly of AT3 occurs via a helical bundle that buries ∼4,200 Å^2^. In solution AT3 is in an equilibrium with dodecamers (AT12) featuring a tetrahedral arrangement of AT3 stabilized by a network of ion pairs between the amino terminus of Met1 and the sidechain carboxylate of Asp7 of adjacent trimers. A transient AT6 intermediate is hypothesized to arise from dimerization of AT3 via formation of two sets of Met1-Asp7 ion pairs and burial of ∼1,100 Å^2^; dimerization of two AT6 assemblies to form AT12 is favored by formation of ten additional sets of ion pairs and burial of another ∼4,600 Å^2^.

AT binds asymmetrically to Trp-loaded TRAP. Although *Bsu* TRAP is predominantly undecameric, both undecamers and dodecamers are observed in mass spectra [1,31–33], and crystallographic studies of *Bsu* AT bound to dodecameric TRAP rings revealed AT_3_ docked on the RNA binding surface of TRAP_12_, via protein-protein surfaces of the zinc binding and helical domains that are not accessible in AT_12_ [34]. Mass spectrometric, solution small angle scattering, and analytical ultracentrifugation experiments [35] are also consistent with a model in which only AT_3_ can compete with RNA for the TRAP-RNA binding site. These findings suggest that the AT_3_ ↔ AT_12_ equilibrium may represent an additional regulatory step.

We found experimentally that preventing formation of the inter-trimer Met1-Asp7 ion pairs favors AT_3_, even at high protein concentrations [26]. Addition of excess zinc during protein recombinant production of AT in *E. coli* results in significant accumulation of formylmethionine AT (*f*AT), presumably due to poisoning of peptide deformylase [36,37]; the resulting immature *f*AT can be chromatographically separated from mature AT under denaturing conditions [26]. Other modification to the N-terminus, such as inclusion of a His-6 tag, similarly prevents AT_12_ formation [38]. This provides the means to experimentally produce homogenous samples of either AT_3_ or AT_12_, to allow characterization of the two oligomers in the absence of confounding effects from their equilibrium.

We have leveraged the ability to selectively produce AT trimers and dodecamers to determine the solution structure of *Bsu* AT_3_ and to examine the pH-and concentration-dependent equilibrium between to AT_3_ and AT_12_ using small angle x-ray scattering (SAXS), sedimentation velocity analytical ultracentrifugation (AUC), and native mass spectrometry (nMS). In addition, backbone amide ^15^N NMR relaxation measurements [39–41] allowed us to study the dynamics of both the trimeric and dodecameric AT forms. These results provide new insights into the structural basis of Anti-TRAP’s homo-oligomeric assembly, and how this process may modulate AT’s function in its regulatory context.

## Results

### Anti-TRAP (AT) adopts trimeric and dodecameric species in solution

#### NMR and native MS reveal two states in slow equilibrium

At pH values near 7, purified AT samples at 10^-3^ M concentrations yield ^15^N HSQC spectra that exhibit doubling of many amide resonances; such doubling arises in cases of chemical heterogeneity or slow conformational exchange between two states. After chromatographically separating mature AT from immature formylated AT (fAT) under denaturing conditions [26], samples of mature AT produced the same signal doubling (Figure 3), indicating that it is not due to chemical heterogeneity. Nano electrospray ionization mass spectrometry under soft ionization conditions, at protein concentrations one hundred times lower than NMR (10^-5^ M), yielded two sets of ions, one corresponding to the mass of AT_3_, and the other, AT_12_ (Figure 3). Adjusting the solution pH to 8 produced a higher proportion of AT_3_ ions, while at pH 6 similar intensities were observed for both sets of ions, consistent with prior findings from pH-dependent NMR spectra. In addition to affecting the oligomeric state, changing the pH slightly altered the preferred charge state distribution: at pH 8, the 7+ and 14+ charge states were favored for AT_3_ and AT_12_, respectively; at pH 6, the 8+ and 15+ charge states dominate the spectrum. Although AT_3_ and AT_12_ populations varied with pH, no other oligomeric species (e.g., AT_1_, AT_6_) were observed in the nMS data over a wide range of conditions.

**Figure 3.**
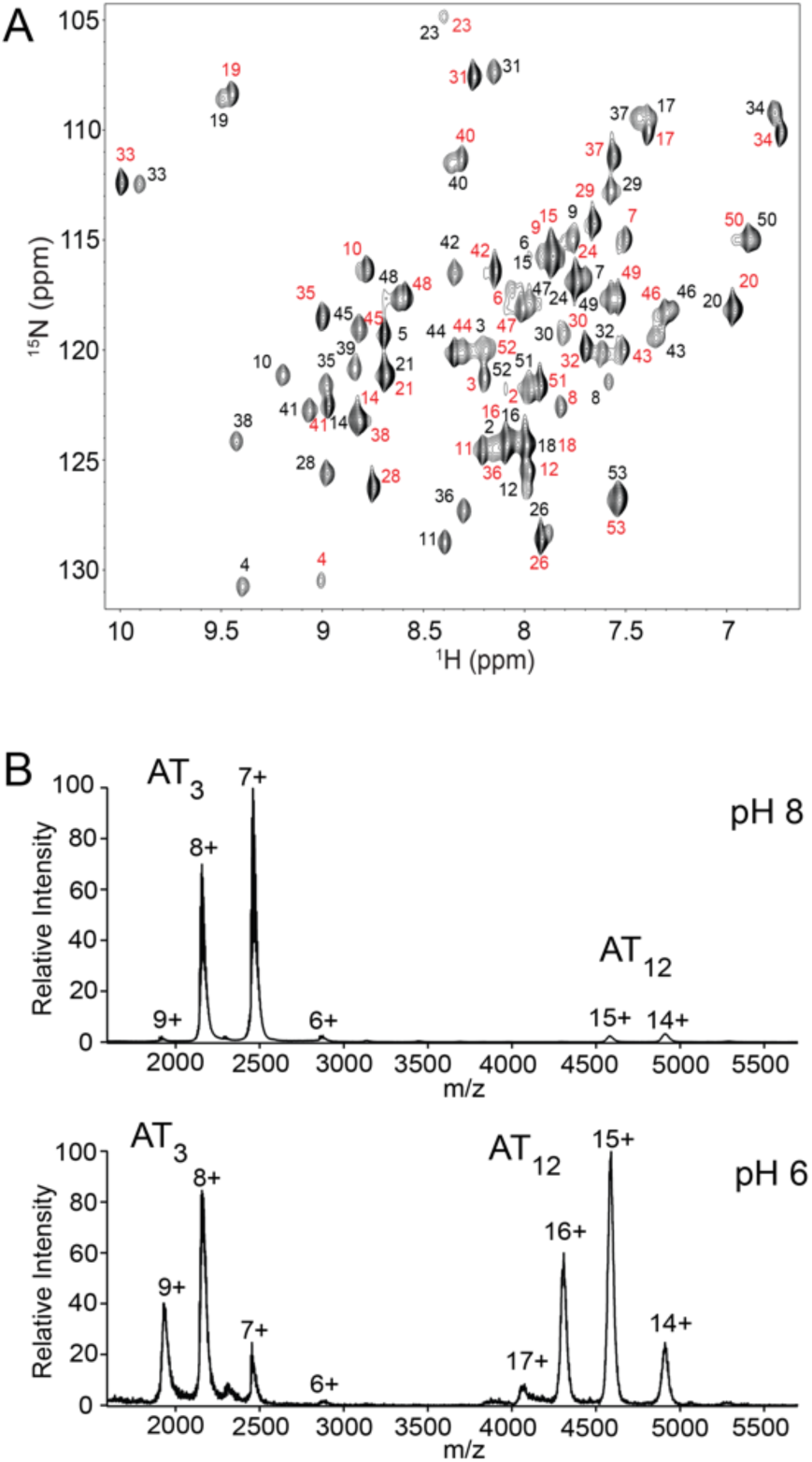
AT adopts trimeric and dodecameric states in solution. (A) ^15^N-^1^H NMR spectrum of 1 mM AT, pH 7.5 reveals doubling of many signals; residue assignments as indicated (black, AT3; red, AT12). (B) Native MS in 100 mM AmAc shows trimers are favored at pH 8 and 12-mers are favored at pH 6; predicted (Zn-AT)3: 17,144.9 Da; (Zn-AT)12: 68,579.4 Da; observed at pH 6: 17,181.8 ± 63.7 Da and 68,826.5 ± 76.4 Da; observed at pH 8: 17,181.5 ± 55.3 Da and 68,751.5 ± 47.3 Da. Other oligomeric states are not evident. The extra mass can be attributed to salt and/or solvent adducts, typical for the instrument used.

#### Sedimentation velocity experiments provide evidence for a lowly populated hexamer intermediate

Guided by prior sedimentation velocity experiments at a fixed pH that revealed both AT_3_ and AT_12_ in solution [23], we performed analytical ultracentrifugation sedimentation velocity experiments at pH 7, 7.25, 7.5, and 8 and a range of AT concentrations (Figure 4). For systems with components undergoing rapid exchange on the timescale of the sedimentation experiment, the modeling of boundary phases can provide the sedimentation coefficients for the components as well as the stepwise binding rates [42]. Sedimentation velocity data at each pH was globally fit across three concentrations in the program SEDPHAT [43], initially to a simple two-state model comprising the two states observed in NMR and MS spectra,

**Figure 4.**
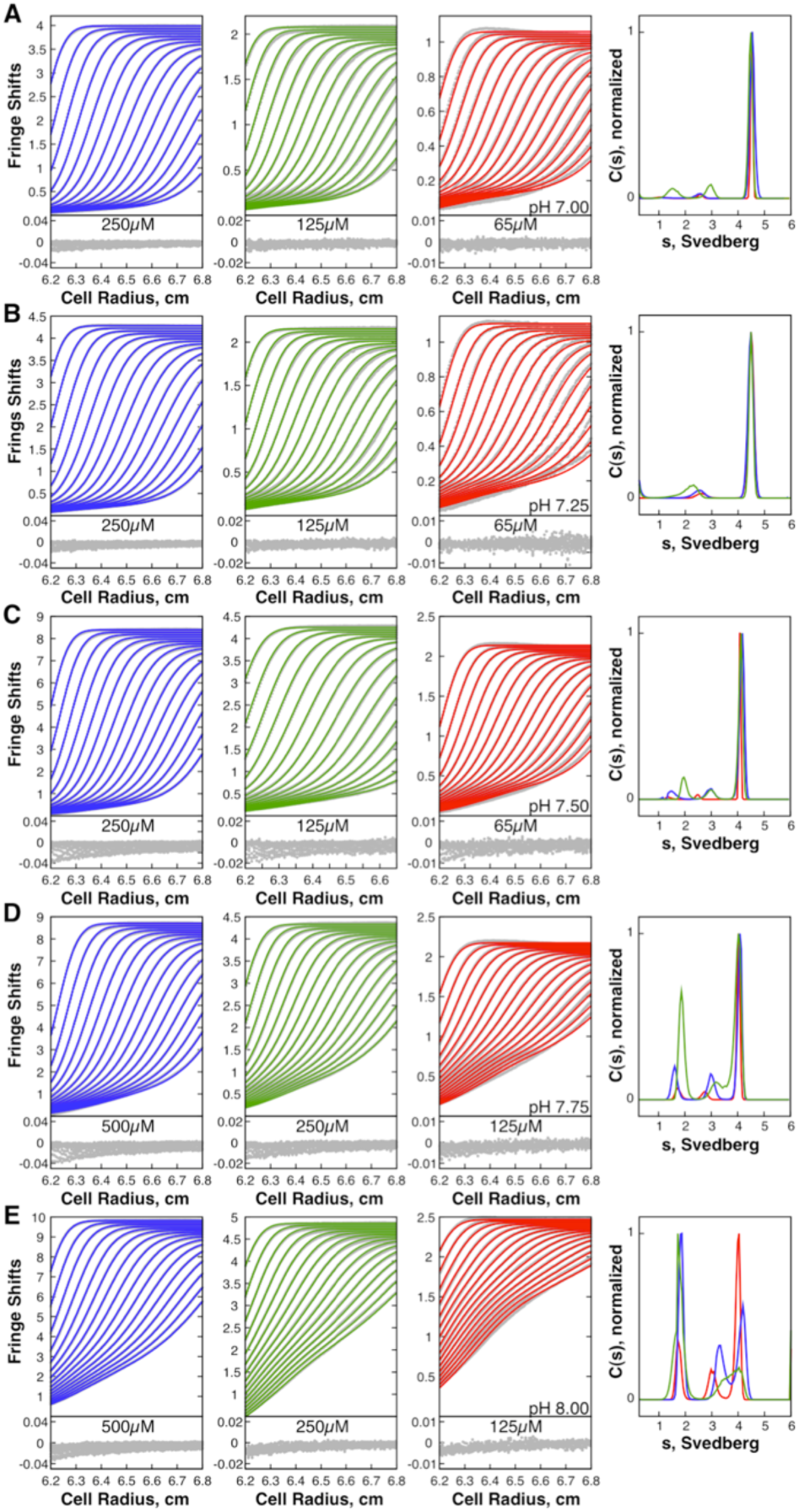
Sedimentation velocity experiments over a range of pH values can be fit with a three-state association model AT3 Η (AT3)2 Η AT12. Interference fringe shifts (gray dots) are globally fit across three AT concentrations (first three columns). Fits provided by the mechanistic model are in blue, green, and red lines. Only every tenth scan is shown. Residuals are below the data. Fourth column, sedimentation coefficient distributions with blue, red and green corresponding to the concentrations on the left, at pH 7.00 **(A)**, pH 7.25 **(B)**, pH 7.50 **(C)**, pH 7.75 **(D)**, and pH 8.00 **(E).**

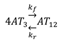

The resulting fits were poor, especially at alkaline conditions, and were better fit with a three-state exchange model,

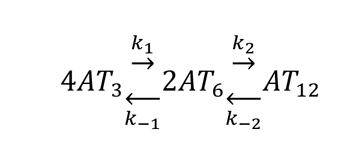

Lamm modeling of the sedimentation data predicts species with an intermediate sedimentation coefficient, which we attribute to a small population of hexamer AT_6_, despite not observing this species by nMS or NMR (Figure 3). Fits to experimental interference fringe shifts using this model permitted optimization of two sets of apparent rate constants describing dimerization of AT_3_ trimers into AT_6_, and a subsequent dimerization of two AT_6_ into AT_12_. From the ratio of the fitted rate constants, we obtained consistent equilibrium dissociation constants for the first step, ranging from 20-90 µM over the experimental pH range. For the second step, the association rate constant dropped over three orders of magnitude over the same pH range, resulting in equilibrium constants from 0.05-50 µM. Thus, at low pH the second step is highly favored compared to the first step, resulting in very low concentrations of the hexameric intermediate AT_6_. This is generally consistent with the relatively low populations of intermediate species in the fitted c(s) distributions (Figure 4). The excellent agreement between experimental and modeled data suggests that monomeric AT is not present at a significant population.

#### SAXS profiles of AT can be reconstructed from those of AT3 and AT12 oligomers

We investigated the pH-dependent structure of AT using small angle x-ray scattering (SAXS). For mixtures, the resulting scattering profiles are the weighted sum of scattering profiles of the individual components [44]. Experimental scattering curves were de-convoluted using form factors for trimeric and dodecameric AT (Figure 5). The predicted scattering profiles for AT_3_ and AT_12_ were calculated with CRYSOL [45] from the trimer and dodecamer assemblies in the crystal structure (2BX9) [28]. Deconvolution of the experimental data from the polydisperse AT solutions was performed using OLIGOMER [44]. The resulting fits using these two components describe the experimental data well, with AT_12_ dominating below pH 7.5. We also tested for the presence of an inverted dodecameric structure such as that observed in crystallographic studies of *B. lichenformis* AT (PDB 3ld0) [29] by including its form factors in fitting the experimental data; this component was found to have zero weight in all of the tested experimental conditions. These data are consistent with a slow exchange between AT_3_ and AT_12_ under near-physiological conditions, and a negligible population of intermediates.

**Figure 5.**
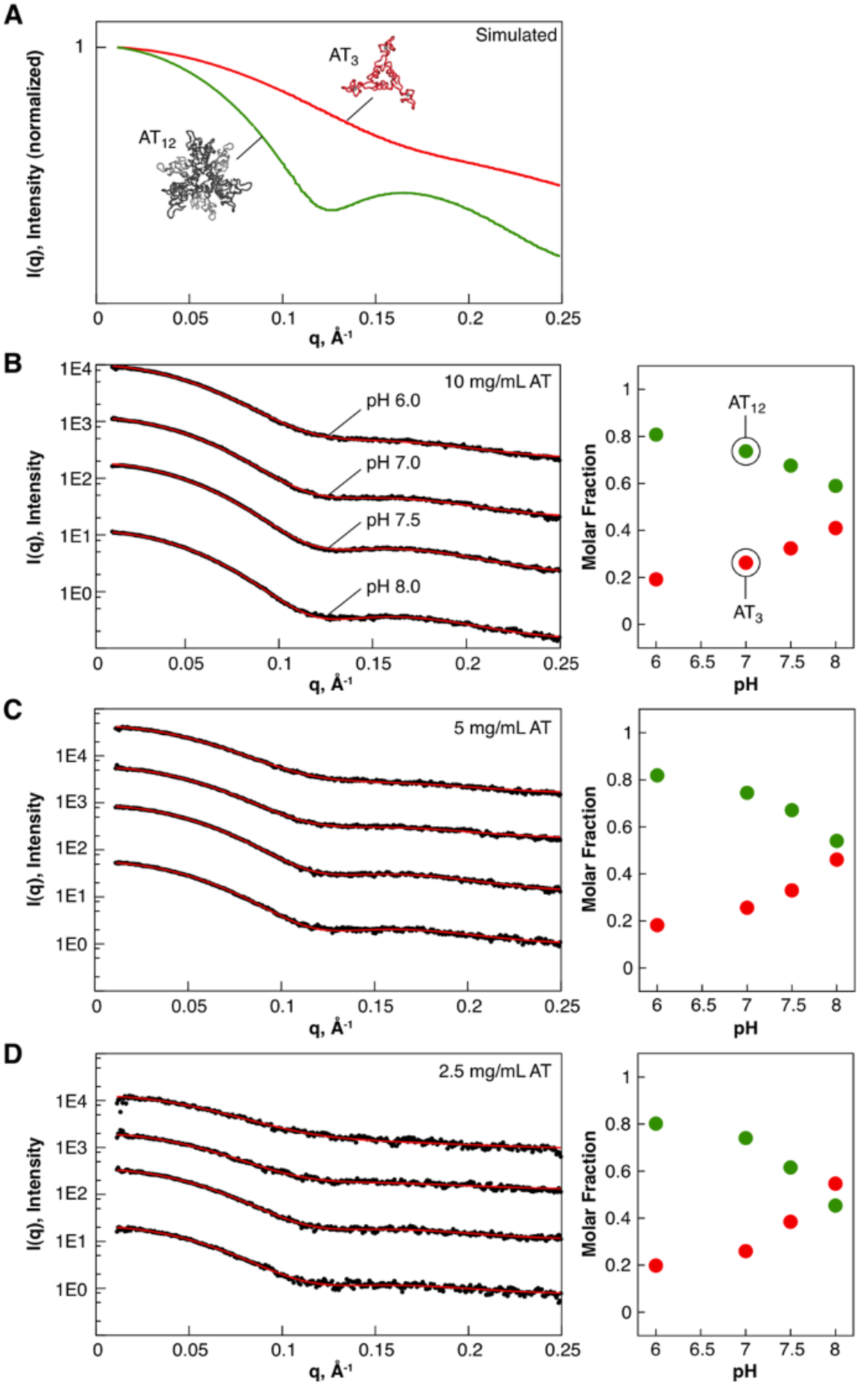
SAXS profiles of AT can be described as mixtures of AT3 and AT12. A, scattering curves for AT3 and AT12, computed from the dodecameric crystal structure (2bx9). B-D, left, experimental scattering intensities (black) for AT recorded over a range of pHs (6, 7, 7.5, 8) and concentrations (1.77, 0.89 and 0.44 mM), fit to a two-state model with varying fractions of AT3 and AT12. Profiles are offset in the y-axis from each other for visibility. Right, fitted volume fractions as a function of pH for each sampled concentration.

### NMR studies of fAT3

#### Formylated AT is trimeric with high quality NMR spectra

We used immature *f*AT for detailed investigation of the structure and dynamics of the trimeric TRAP-binding oligomeric form. Since an ion pair between the protonated amine of Met1 and the sidechain carboxylate of Asp7 is proposed to stabilize the dodecameric structure [26,28], the formyl group prevents formation of the ion pair and *f*AT trimers do not oligomerize to form detectable dodecamers, even at millimolar concentrations at low pH. Resonance assignments and NMR structural data were recorded at 55 °C, to match the conditions used for backbone resonance assignments of *Bst* TRAP [26,46,47].

NMR spectra of *f*AT_3_ are of exceptionally high quality (e.g., Figure 6), enabling nearly complete backbone and side chain resonance assignments. *Bsu* AT comprises 53 amino acids, and zinc-bound *f*AT_3_ has a nominal molecular mass of 17.1 kDa. Three-fold symmetry and fast tumbling result in narrow NMR resonances and high-quality heteronuclear correlation spectra. Backbone resonances were assigned from standard triple resonance spectra, while sidechain assignments were obtained from a combination of 2D and 3D double-and triple-resonance spectra. This allowed for assignment of every proton in the protein except for the side chain amine and hydroxyl protons.

**Figure 6.**
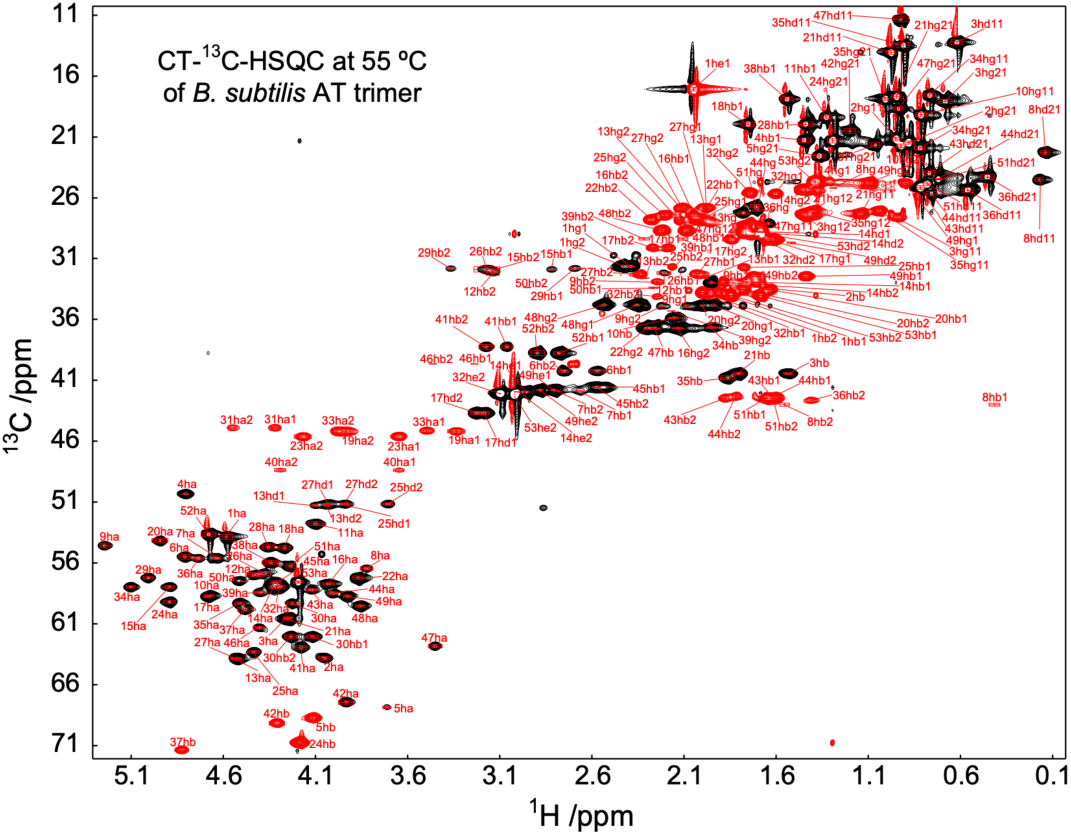
Symmetry and exceptional NMR spectra of *f*AT3 allowed nearly complete resonance assignments. Aliphatic region of a two-dimensional constant-time ^1^H-^13^C correlation spectrum of *f*AT3. In this spectrum ^13^C resonances coupled to even (red) and odd (black) numbers of aliphatic carbons have opposite sign. Aliphatic carbon resonance assignments as indicated.

### The structure of fAT3 is unperturbed by pH

SAXS, nMS and NMR data demonstrate that the trimer-dodecamer equilibrium is strongly affected by pH (Figure 3, Figure 4, Figure 5). This behavior was attributed to a near-neutral pKa for the amine at the N-terminus of the protein [30], and its ion pair with the carboxylate side chain of Asp7 [26,28]. There is one other functional group in AT with a predicted pKa near neutrality: His50, which is conserved among AT variants [19,29]. In crystals the side chain imidazole Nε2 of His50, located at the end of the C-terminal helix, is observed to form an intra-chain ion pair with the carboxylate of Glu9, located in the zinc-ribbon motif (Figure 2). Changes in the protonation state of the imidazole side chain could conceivably alter the geometry between the zinc-binding and helical domains and thus alter trimer packing in the dodecamer, driving changes in oligomeric structure (e.g., such behavior has been engineered into homo-oligomeric assemblies [48]). To examine whether His50 or another titratable group in the protein might induce structural changes that contribute to the pH-dependent oligomerization behavior, we recorded ^1^H-^15^N correlated NMR spectra of *f*AT over a range of pH values (Figure 7). Since *f*AT has a blocked N-terminus and does not form stable dodecamers, this allowed distinguishing spectral effects of ionization from those resulting from oligomerization. Correlation spectra of *f*AT recorded over a pH range that would shift the predominant oligomeric state from AT_3_ to AT_12_ in the mature protein. result in only minor shifts in the spectra, particularly the amides of Lys49 and Leu51, reflecting their proximity to the His50 indole. This is consistent with a dominant role of a near neutral pKa for the N-terminal amide group in stabilizing the dodecamer.

### Solution Structure Determination

The solution structure of the TRAP-binding oligomeric state of *Bsu* AT (AT_3_) might be inferred from the 2.8 Å crystal structure of the dodecamer (2BX9)[28], the 2.1 Å structure of the *Bli* dodecamer (3LCZ)[29], and 3.2 Å structure of the *Bsu* AT bound to *Bst* TRAP (2ZP8) [34], but *de novo* structure determination was justified by the potential for perturbations from oligomerization and crystal packing. To that end, we determined the solution structure of *f*AT_3_ using distance restraints from NOEs, orientation restraints from residual dipolar couplings, dihedral angle restraints from three-bond scalar couplings, three-fold symmetry restraints, and overall shape restraints from SAXS scattering curves.

Because of its homo-oligomeric structure and diverse source of restraints, structure determination proceeded via several steps. First, three-dimensional models of the zinc-bound AT protomer were computed from torsion angle restraints and NOEs that could be unambiguously assigned as being intramolecular. Next, we determined the solution structure of the trimer by replicating the chain and assigning as ambiguous all NOEs for which conservative distance filters prevented their assignment as being intramolecular. Most ambiguous NOEs could be assigned via iterative structure refinement steps, though some NOEs remained ambiguous throughout the refinement process (Table 1).

**Table 1:**
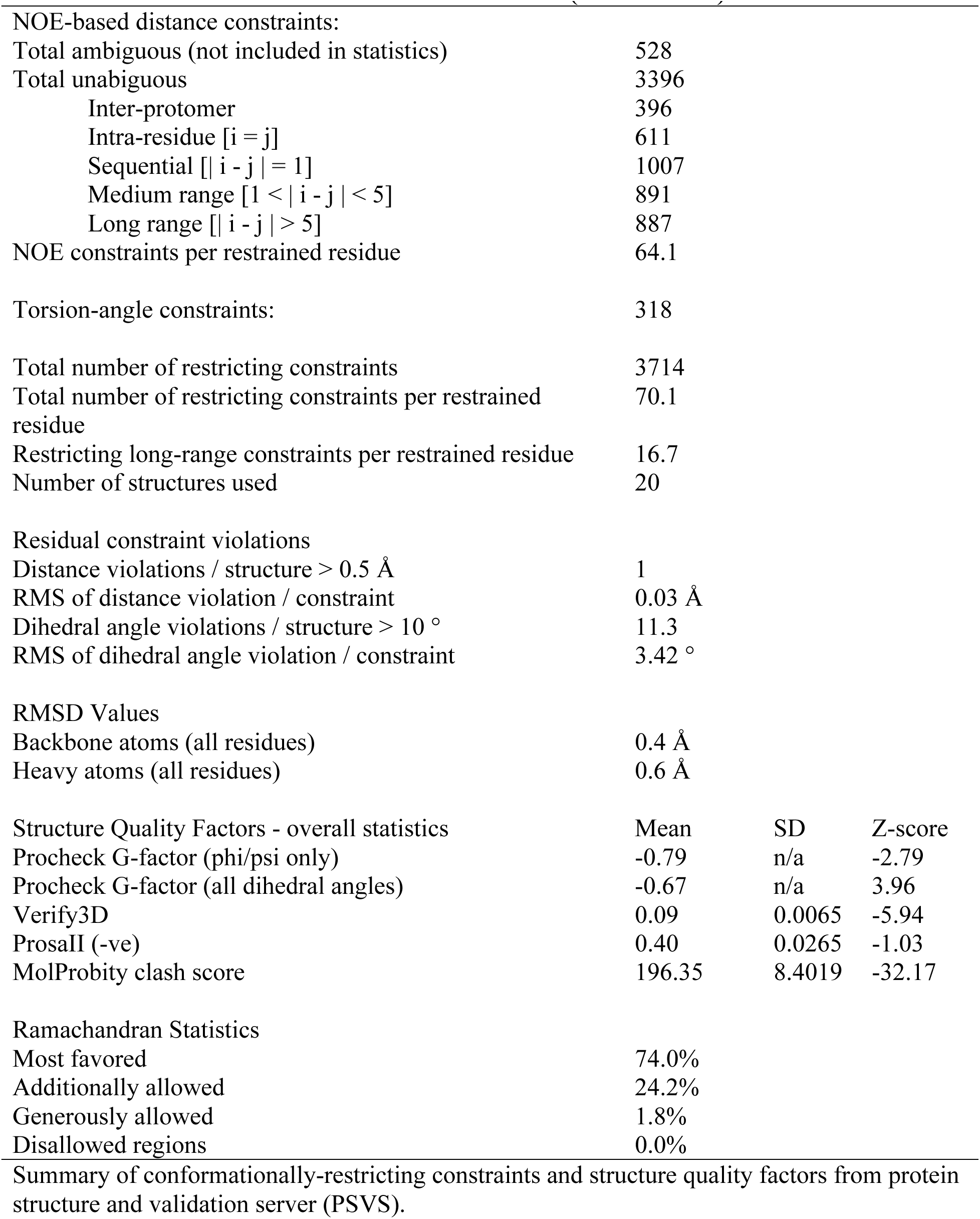
NMR structural statistics for AT ensemble (20 structures):

**Figure 7.**
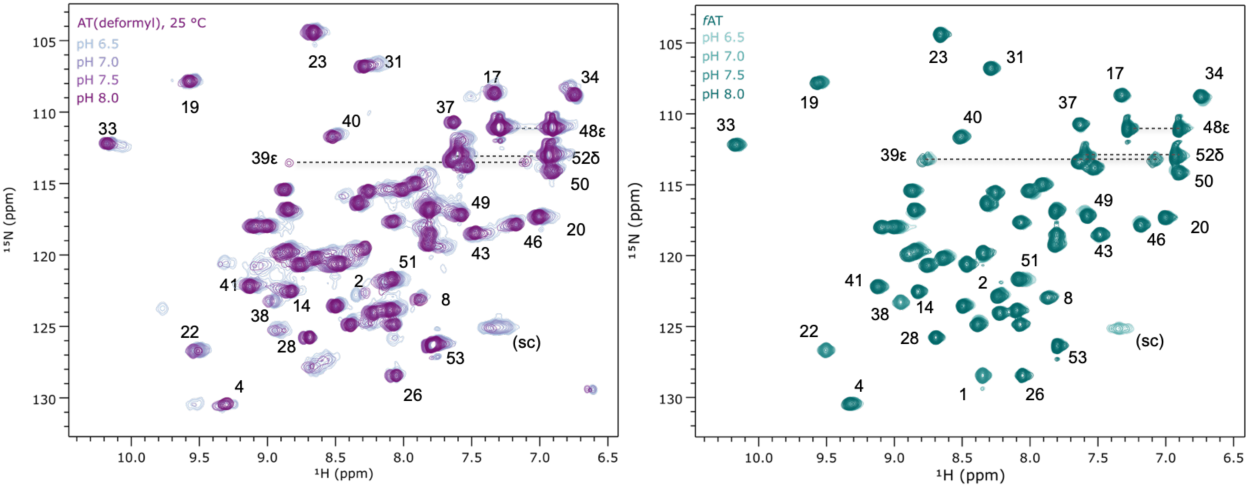
Protonation of the Met1 amine is coupled to oligomerization of AT. Left, ^1^H-^15^N correlated spectra of AT recorded at 25 °C and pH 6.5, 7.0, 7.5 and 8.0 reveal strong perturbations of several signals, and line narrowing at high pH, which favors trimeric AT. Right, for *f*AT the N-terminal amine is protected by the formyl group and very minor shift perturbations are observed only for residues flanking His50; this indicates no pH-dependent structural shifts in the AT trimer. Amide assignments transferred from 55 °C via a temperature series. (sc), side chain amide.

Symmetry, residual dipolar coupling (RDC) and SAXS restraints significantly increased the precision of the NMR structure (Figure 8). Ensembles computed using only NMR-derived distance and torsion angle restraints yielded poor convergence, with an overall backbone RMSD of 1.5 Å, and per-protomer backbone RMSD of 1.3 Å. RDCs and SAXS data improved the overall precision, but not as much as inclusion of three-fold symmetry restraints, which increased the overall precision by almost an Angstrom (from 1.5 Å to 0.6 Å).

**Figure 8.**
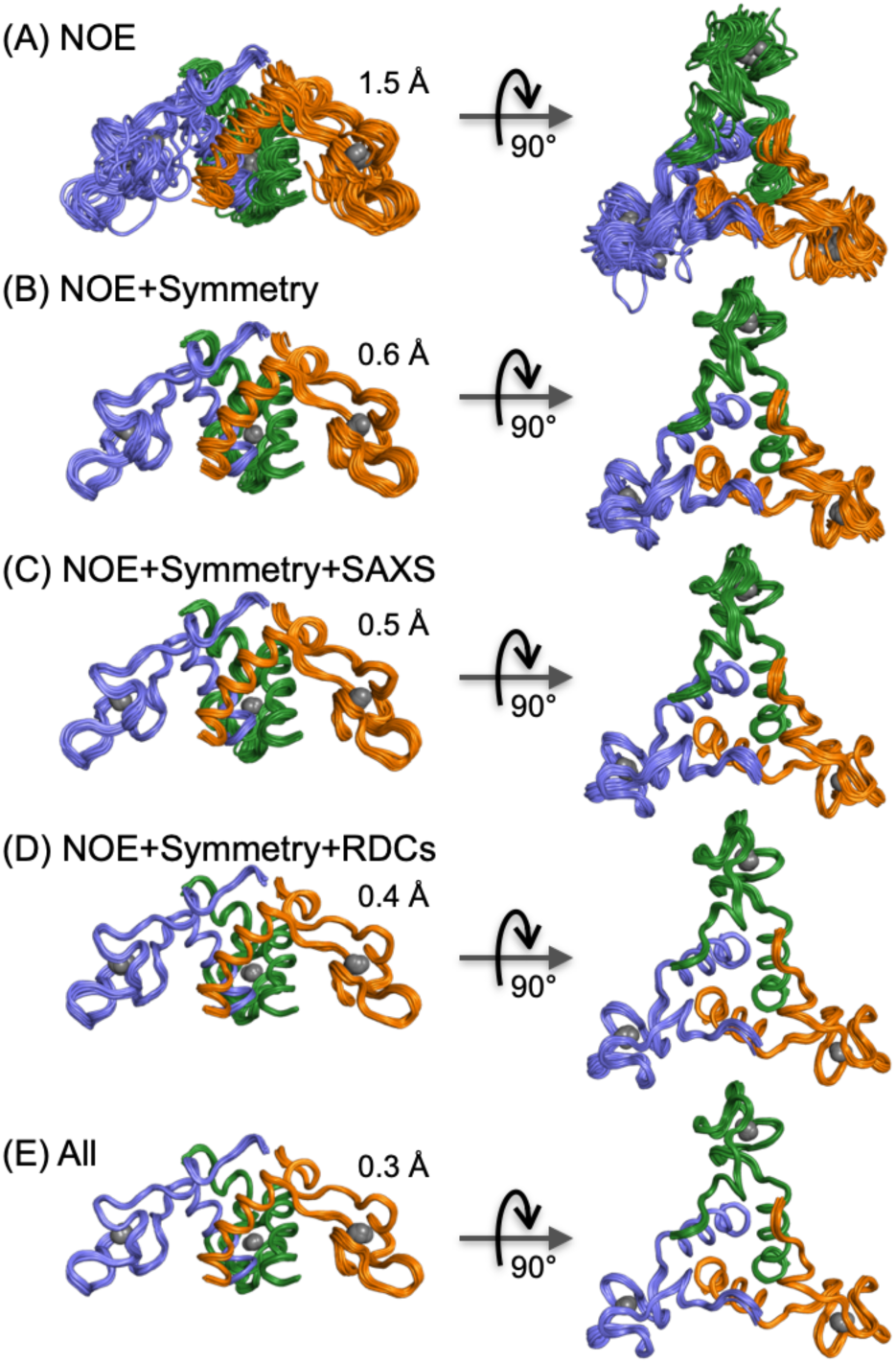
Impact of restraint classes on precision of AT3 NMR ensemble. Each ensemble was computed using NMR-derived torsion angle restraints, and (A) NOEs, (B) NOEs and three-fold symmetry restraints, (C) NOEs, symmetry, special restraints from SAXS data, (D) NOEs, symmetry restraints and residual dipolar couplings, and (E) all restraint classes. Backbone RMSDs as indicated.

### Comparison to crystallographic and AlphaFold2 models

The final NMR ensemble was well defined by the restraints, with a backbone RMSD of 0.3 Å for C^α^ atoms (Figure 8, Figure 10, Table 1), and reveals a high degree of similarity with the trimers in the *B. subtilis* dodecamer (RMSD ∼0.6 Å; Figure 10). This variance is similar to that observed between protomers in the experimentally determined crystal structure (∼0.6 Å for backbone atoms in 2BX9), and within the variance between those structures and that predicted by AlphaFold2 [49]. These subtle conformational differences imply intrinsic flexibility in the protein.

**Figure 9.**
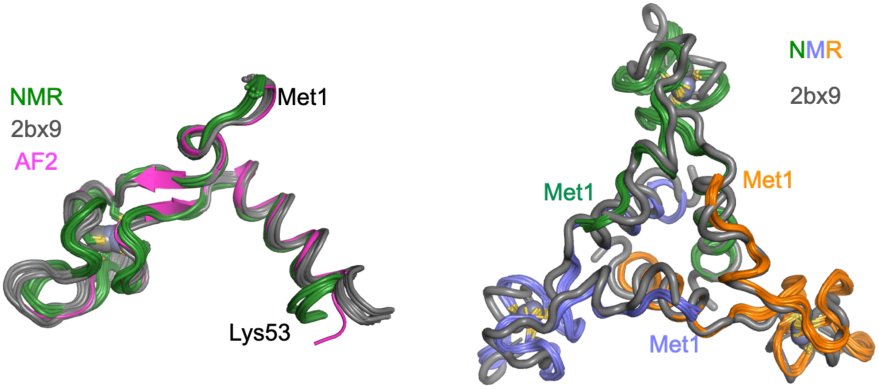
Amide ^15^N relaxation rates for trimeric and dodecameric AT reveal differences in overall correlation and local dynamics. Relaxation rates were measured at 600 MHz, 55°C. Local correlation times were computed from the *R*2/*R*1 ratios {Palmer}. Secondary structures and location of metal ligands are indicated schematically above.

**Figure 10.**
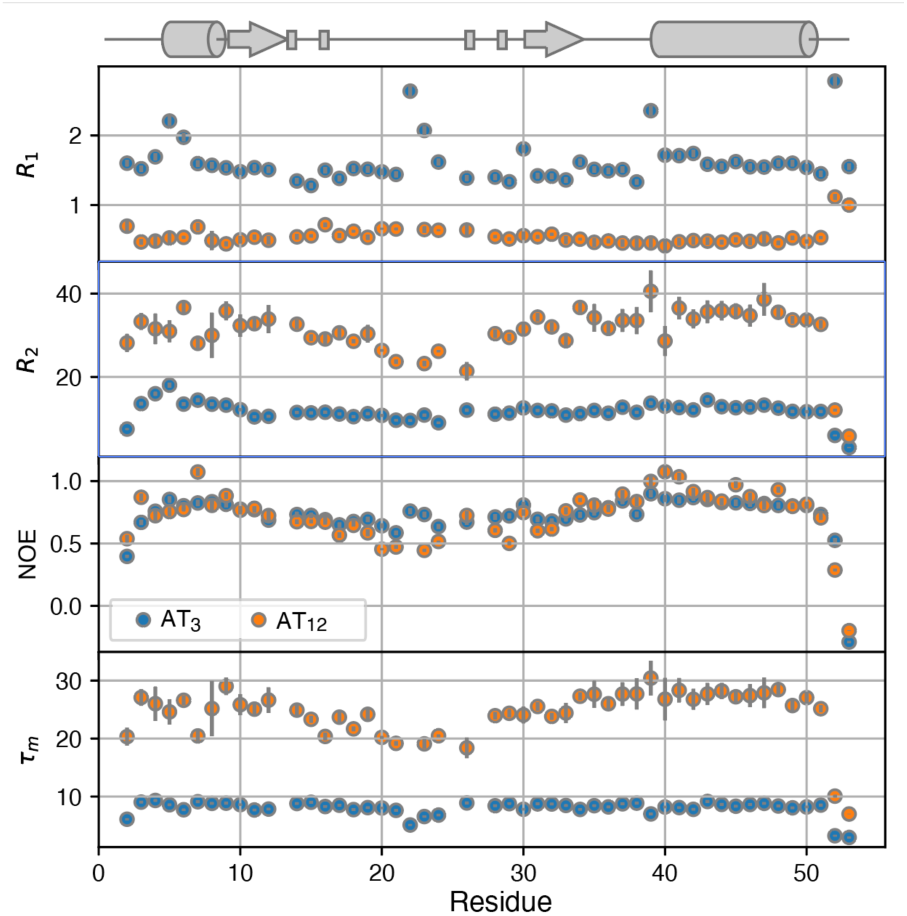
Solution structure of AT is congruent with its structure in AT12 and that predicted by AlphaFold2. Left, green, NMR ensemble of 20 conformers for chain A; grey, superposition of the twelve chains in the 2BX9 crystal structure; pink, AlphaFold2 prediction. Right, superposition of the AT3 NMR derived ensemble with chains A-C (blue, orange, green) in the 2BX9 crystal structure.

### AT dynamics

Backbone amide ^15^N *R*_1_, *R*_2_, and ^1^H-^15^N NOE relaxation rates were measured on both the 68 kDa dodecameric AT_12_ and 17 kDa trimeric AT_3_ using conventional two-dimensional heteronuclear correlation spectra (Figure 9) [40,41]. Differences in the *R*_1_ and *R*_2_ relaxation rates for the two oligomeric states reflected the large difference in their rotational diffusion tensors; overall correlation times τ_m_ computed from the 10% trimmed mean *R*_2_/*R*_1_ ratios were 8 and 25 ns at 55 °C, respectively [50]. Hydrodynamic calculations based on the crystal structure of AT_12_, and the *R*_2_/*R*_1_ ratios were also consistent with an isotropic diffusion tensor and an overall τ_m_ of 25 ns. However, *R*_2_/*R*_1_ ratios for AT_3_ were consistent with an axially symmetric diffusion model with an anisotropic diffusion tensor, D_iso_ = 2.00 ± 0.01 x 10^7^ s^-1^, *D*_║/┴_ = 0.82 ± 0.04 (χ_m_ = 8.3 ns).

Large differences in the rotational diffusion tensors of AT_12_ and AT_3_ are accompanied with differences in internal motions. Amide ^15^N relaxation rates were analyzed using the extended model-free formalism [51–53] to compare the amplitude internal dynamics of AT_3_ and AT_12_ on the ps-ns timescale, as represented by the square of the generalized order parameter, *S^2^* (Figure 11). While both proteins exhibited generally high-order parameters, AT_3_ and AT_12_ were found to exhibit local differences in fast internal motions. High *S*^2^ values for AT_3_ imply rigidity of the C-terminal helical bundle, and larger amplitude fast motions in the zinc binding arm domains and the N-terminal helix. By contrast, the zinc binding domains in AT_12_ are relatively rigid on the ps-ns time scales reported by *S*^2^, likely a reflection of their packing onto adjacent zinc binding domains in the dodecamer (Figure 2). The two oligomeric forms of AT also exhibit different patterns of the phenomenological exchange broadening term *R*_ex_, attributable to dynamics on slower µs-ms time scales [53,54]. For AT_3_, the residues with largest *R*_ex_ terms are located at the N-terminus of the protomers, with smaller *R*_ex_ terms at the zinc binding domains. For AT _12_, almost the inverse is observed, with the largest exchange terms needed to fit data for the helical bundle.

**Figure 11.**
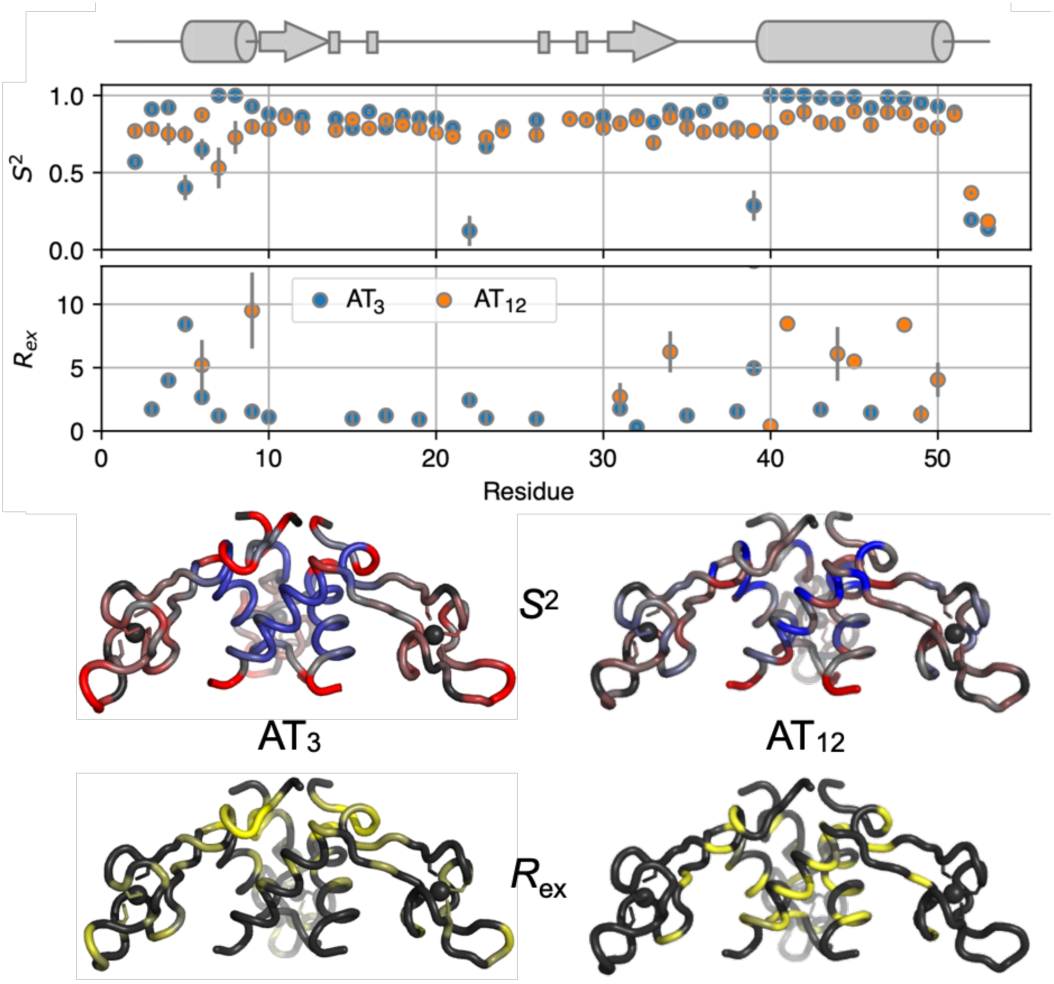
Fitted Model-Free dynamics parameters suggest a stable helical bundle and flexible zinc-binding domains in AT3, but less variable dynamics in AT12. Top, fitted model free order parameters *S*^2^ and *R*ex for AT3 and AT12. Middle, fitted *S*^2^ values mapped to the backbone of the AT trimer using a linear color ramp from the mean value (grey) to + (blue) or – (red) two standard deviations from the mean. Bottom, *R*ex values mapped to the AT3 structure with a color ramp from 0 to ≥ 4/s (yellow).

## Discussion

Tryptophan biosynthesis is highly regulated in bacteria [18,55,56]. Within bacteria, TRAP-mediated transcription attenuation and translation repression via RNA binding represents an unusual regulatory mechanism. In the subset of organisms that also encode Anti-TRAP, its expression represents yet an additional regulatory layer by binding to and reversing the activity of an inhibitory protein. In this work we examined the equilibrium between active and inactive oligomeric states of *Bsu* Anti-TRAP using nMS, SAXS, AUC and NMR spectroscopy. We determined the solution structure of the active AT_3_ form of the protein and observed differences in its internal dynamics relative to that of the inactive form AT_12_. These studies thus help to inform our understanding of how AT inhibits RNA binding by Trp-bound TRAP, and of how the AT_3_ ↔ AT_12_ equilibrium may provide yet another regulatory layer.

In our studies of *Bsu* AT, serendipity played a significant role. The fortuitous requirement of a mature amino-methionine terminus for stabilizing the AT dodecamer coincided with experimental zinc supplementation of minimal media for recombinant protein expression, which likely prevented maturation of a sizeable fraction of the protein. The second residue in *Bsu* AT is a valine, which is often a poor substrate of methionine aminopeptidase [57,58], thus ensuring retention of Met1 in the mature protein. The application of NMR and MS enabled detection of the formylated N-terminus, while denaturing reversed-phase chromatography allowed separation of immature *f*AT from mature AT, which was not possible with conventional native purification. While the AT_3_ ↔ AT_12_ equilibrium is evident in samples of mature AT, the ability to prepare complexes of pure *f*AT provided an unambiguous means of establishing the role of the N-terminal amine in the equilibrium and enabling independent structural characterization of the TRAP-binding form of the protein (AT_3_).

We found that *Bsu f*AT_3_ (and AT_12_) yields NMR spectra of exceptional quality, enabling complete resonance assignments and high-resolution structure determination by application of NMR, symmetry and SAXS restraints. Symmetry restraints were found to strongly increase the precision of the structural ensemble, while SAXS and RDC restraints provided modest improvements. We found the solution structure of AT_3_ to be indistinguishable from that in the 12-mer crystal, or from the AF2 predicted model, indicating little conformational adjustment upon formation of the dodecamer.

While the average structure of the AT trimer in isolation and in the dodecamer are highly similar, they differ in their dynamics. Consistent with the overall size and shape of the oligomers, backbone ^15^N relaxation measurements revealed an isotropic rotational diffusion tensor for AT_12_ and axially symmetric diffusion for the smaller AT_3_. Model-free analysis of internal dynamics revealed flexible zinc ribbon motifs in AT_3_, suggesting a role for the dynamics of these motifs in engaging Trp-bound TRAP rings. AT_12_ has comparatively more rigid zinc-binding domains, likely reflecting packing constraints from the other trimers in the tetrahedral arrangement. Modest *R*_ex_ terms in the N-terminal and zinc-binding domains of AT_3_ may reflect transient but unproductive AT_3_-AT_3_ association reactions, whereas the pattern of *R*_ex_ terms is different in AT_12_, perhaps reflecting transient dissociation to the AT_6_ intermediate.

We conclude that the *Bsu* AT_3_-AT_12_ equilibrium is dependent on the near-physiological pKa of the Met1 amine, with AT_12_ favored by a protonated N-terminus and AT_3_ favored when unprotonated. Prior NMR observations [26] are corroborated here by nMS, SAXS, and AUC experiments as a function of pH. NMR spectra of *f*AT, over a pH range that alters the trimer-dodecamer equilibrium for mature AT, only show minor shift perturbations adjacent to His50, supporting the dominant role of the protonated N-terminus in stabilizing the dodecamer (Figure 7). Native MS spectra (at lower protein concentration) reveal AT_3_ as the major species at pH 8, while at pH 6 nearly equal intensity is observed for ions arising from AT_3_ and AT_12_ species (Figure 3); this provides some context to a prior nMS study which found exclusively AT_12_ when sprayed from pH-unadjusted ammonium acetate [25,34], which can approach a value of 4.75 in the ESI plume [59]. SAXS data for mature AT recorded as a function of pH and protein concentration could be well described by considering varying ratios of AT_3_ and AT_12_ (Figure 5). Experimental curves could be reproduced form a linear combination of the form factors computed from the crystallographic dodecamer and a trimer extracted from the dodecamer. This finding is consistent with the failure to observe other assembly intermediates under equilibrium conditions. Sedimentation velocity (AUC) experiments further corroborate the pH-dependent equilibrium between AT_3_ and AT_12_ and provide evidence for a short-lived AT_6_ intermediate. If we consider AT_3_ a “monomer” that can tetramerize, this behavior is consistent with biophysical characterization of other “tetramers”, which predominantly follow a monomer-dimer-tetramer oligomerization path [60]. Small surface burial and minimal polar contacts in AT_6_, result in transient/weak dimerization; large surface burial and many polar contacts favor AT_12_ upon collision of two AT_6_ “dimers of trimers”.

## Conclusion

The mechanism by which Anti-TRAP (AT) promotes Trp production by inhibiting RNA binding by tryptophan-activated TRAP remains poorly understood [19,23,34,35]. The capacity of AT to form stable dodecameric and trimeric structures, wherein only the trimeric form is competent for binding and inhibiting TRAP is suggestive of a regulatory role for that equilibrium. While the complementary biophysical studies reported here provide qualitatively consistent insights into the properties of AT and its mechanism of oligomerization and TRAP inhibition, further work would be required to establish quantitative consistency. Some sources of uncertainty include the effect of molecular tumbling on observed NMR signal intensities, of ionizable buffers and of protein charge on gas phase ionization and gas phase signal detection, and relatively low structural resolution of SAXS and sedimentation velocity methods. Those caveats aside, it is clear that in vitro AT exhibits a slow equilibrium between active AT_3_ and inactive AT_12_ states.

This slow association/dissociation behavior of AT could serve a regulatory function by kinetically modulating the response to changes in Trp concentration [61]. In this model (Figure 1), TRAP is slow to release Trp upon decrease in its cellular concentration and remains active for RNA binding and inhibiting *trp* expression. Meanwhile, decreasing cellular Trp results in accumulation of uncharged tRNA^Trp^ and T-box mediated activation of AT expression. Expressed AT will trimerize and rapidly bind to RNA-free Trp-activated TRAP rings, preventing them from binding new *trp* RNA leaders. AT_3_ that don’t find TRAP rings could instead dodecamerize, resulting in slowly equilibrating populations of active AT_3_ and inactive AT_12_. This scenario would thus generate a pool of fast-acting AT_3_ and a reservoir of slowly activated AT_12_. The resulting steady-state equilibrium between AT_3_, AT_12_, AT_3_-Trp-TRAP_11_, Trp-TRAP_11_ and Trp-TRAP_11_-RNA would achieve a steady state of Trp homeostasis.

The properties of the AT_3_ “triskelion”, with its polydentate configuration of flexible zinc-binding arms, seem uniquely suited for capturing one or multiple TRAP rings, and thereby preventing RNA wrapping. The use of the amino terminus to mediate oligomerization represents a uniquely tuned equilibrium since the pKa and association constants are coupled, such that high protein concentration would shift the pKa to favor the protonated state, and thus the oligomer. It remains unclear why some species of Bacilli evolved to produce AT as an extra layer of regulation (e.g., *Bsu*, *Bli*), while others did not (e.g., *Bst, Bha*). This study highlights the lengths to which organisms have evolved sophisticated mechanisms for metabolic control, and the power of complementary biophysical measurements to describe those properties that enable regulated function.

## Materials and Methods

### Growth and purification of B. subtilis Anti-TRAP

AT was recombinantly expressed from *E. coli* from an engineered codon-optimized expression vector [26]. *B. subtilis* Anti-TRAP is a 53 amino acid polypeptide, with the sequence: MVIATDDLEV ACPKCERAGE IEGTPCPACS GKGVILTAQG YTLLDFIQKH LNK. It has a molecular weight of 5649.57 Da and its predicted extinction coefficient at 280 nm is 1490 M^-1^ cm^-1^ in its reduced form (Expasy ProtParam). Uniformly N-terminal formylmethionine Anti-TRAP (ƒAT) and deformylated AT were purified as previously described [26]. Briefly, transformed BL21(DE3) cells grown and induced in modified M9 minimal media were lysed and cellular debris pelleted by centrifugation. Isotope labeling was achieved by replacing unlabeled ammonium chloride with ^15^N ammonium chloride for ^15^N labeling, by substitution of ^13^C glucose for ^13^C/^15^N double labeling. Heating of the cell lysate supernatant for 15 minutes at 70°C and subsequent centrifugation removed most native *E. coli* proteins. The resulting supernatant was loaded onto Q-sepharose beads (GE healthcare) and eluted via a NaCl gradient. Eluted Anti-TRAP was denatured by addition of 6 M guanidinium and subjected to reverse-phase HPLC purification. AT (i.e., deformyl) and ƒAT elute at sufficiently resolved retention times from a C4 column to be separated and lyophilized, providing >99% desired species. Subsequent refolding results in intact ƒAT trimers for ƒAT, and AT_3_ + AT_12_ in equilibrium for (deformylated) AT.

### Mass spectrometry

The theoretical mass of AT (from ExPASy ProtParam): Monomer (without Zn) = 5,649.57 Da; monomer with Zn (65.38 Da) = 5714.95 Da; trimer with three Zn = 17,144.85 Da; dodecamer with twelve Zn = 68,579.40 Da. AT samples were buffer exchanged twice into 100 mM ammonium acetate at 3 different pH’s (pH 6.8, unadjusted; pH 8, adjusted using ammonium hydroxide; pH 6, adjusted using acetic acid) using size-exclusion spin columns (MicroBioSpin-6; Bio-Rad); the final concentration of AT was 10 μM. Borosilicate glass capillary emitters were pulled in-house with a Sutter Flaming/Brown Micropipette puller P97; samples were loaded and sprayed using nanoESI. Mass spectra were recorded on a Waters Synapt G2S mass spectrometer modified in-house with a SID device located between the trap cell and the drift cell. Data were recorded with capillary voltage, 0.7-1.0 kV; cone voltage, 20 V; extraction cone, 5-8 V; helium gas flow, 120 mL/min; trap gas flow, 2 mL/min; ion mobility nitrogen gas flow, 60 mL/min; source temperature 20 °C; trap wave velocity, 160 m/s; trap wave height, 4 V; ion mobility wave velocity, 300 m/s; ion mobility wave height, 20 V; transfer wave velocity, 65 m/s; transfer wave height, 2 V; time-of-flight analyzer pressure, ∼ 6×10^−7^ mbar; and backing pressure, ∼ 2.5-3 mbar.

### NMR assignments of AT12

AT_12_ predominates at high protein concentration, pH 7 for the mature, deformylated protein. Two dimensional ^1^H-^15^N TROSY spectra of AT_12_ have approximately the number of peaks expected for the protomer, consistent with its molecular symmetry. HNCA, HNCOCA, HNCACB, and HNCO triple resonance spectra recorded at 328 K using water flip-back solvent suppression, sensitivity enhancement, and transverse relaxation optimized spectroscopy (TROSY) coherence selection [62–66] on uniformly doubly labeled (^13^C, ^15^N) AT, allowed assignment of every backbone resonance in the spectrum, corresponding to 48 of the 50 non-proline residues (only M1 and E22 were missing).

### NMR assignments and structural restraints of AT3

*f*AT spectra exhibit signals exclusively attributable to trimeric AT. The NMR spectra of *f*AT3 also exhibit the number of peaks expected for the monomer, consistent with its molecular symmetry. Assignments of uniformly, doubly labeled (^13^C, ^15^N) trimeric AT were obtained using HNCO, HNCACB, CCONH-TOCSY, HCCONH-TOCSY, HCCH-COSY, HCCH-TOCSY, ^15^N-edited TOCSY, constant-time ^13^C-edited HSQC, ^13^C-edited HSQC, constant-time ^13^C-edited HSQC of the aromatic region, ^13^C-edited HSQC of the aromatic region, and a ^1^H-^1^H plane and _1_H-^13^C plane of the HCCH-COSY of the aromatic region [67].

Torsion angle restraints for the backbone φ angles were obtained from the analysis of 3D HNHA spectra [68] and additional backbone torsion angle φ, ψ restraints were obtained from chemical shifts using TALOS [69]. Sidechain ξ_1_ torsion angle restraints were obtained from the HNHB and HACAHB-COSY experiments [68]. The X-Pro amide bonds for AT_3_ were determined to be in the trans conformation from analysis of the Cý and Cψ chemical shifts [70]; Cψ shifts were not assigned for AT_12_ but could be assigned as such in the crystal structure [28]. One bond ^1^D_HN-N_ and ^1^D_N-C_ residual dipolar coupling (RDC) restraints were obtained from IPAP spectra [71,72], of ^13^C/^15^N-labeled AT aligned with 20 mg/ml Pf1 phage (ASLA Biotech). NOE restraints were obtained from a ^13^C/^15^N-edited time-shared NOESY [73] in H_2_O and a ^13^C-edited NOESY [67] in ^2^H_2_O.

### NOE assignments and initial AT3 structure determination

NOE assignments of the symmetrical homo-trimer were made using the symmetry ambiguous distance restraint (symmetry ADR) method starting with a well-defined monomer [74,75]. The structure of the protomers were first obtained by assigning NOEs presumed to be intramolecular, (i.e., NOEs between zinc-coordinating coordinating cysteines, secondary structural elements inferred from secondary structure predictions based on sequence and chemical shift data), and using these as input for restrained torsion-angle refinement with CNS [76]. The set of 20 lowest energy structures (of 35 total) was then used with a 10 Å cutoff to filter through the remaining NOE data. At this stage, NOEs with less than six ambiguous assignments were used for another round of structure refinement and the cycle was iterated while decreasing the distance cutoff by 2 Å until the cutoff was 6 Å and the monomer structure converged to an RMSD of 2.08 Å for backbone atoms. The trimer structure was then calculated by ambiguously assigning any NOEs that could not be satisfied intramolecularly (i.e., they were defined as being between protomers A, B and C) and a structure calculation was performed utilizing non-crystallographic symmetry restraints in addition to the distance, orientation, and torsion angle restraints. This set of structures was then used with a 6 Å filter to assign the remaining NOEs and sort out the interprotomer contacts.

### SAXS and RDC refinement of AT3

To obtain a scattering profile for the trimer, pure ƒAT was purified, refolded and dialyzed against 50mM sodium-phosphate, pH 8, 100 mM NaCl, 200µM ZnCl_2_ and 1mM TCEP. SAXS profiles at 1, 2, 3.3, 5, and 10mg/mL ƒAT (monomer) were acquired at the SIBYLS beamline at the Advanced Light Source in Berkeley, California. Data were obtained at a Q spacing range of 0.01 to 0.32 Å-1, and at 0.5, 1, and 6 second exposure times. The scattering contribution from the buffer was removed by subtracting the scattering profile of the dialysis solution. Processing, validation, and analysis of SAXS data was performed with the ATSAS package [77] and custom scripts. For restraints to the AT_3_ structure, the 6 s exposure profile of the 3.3 mg/mL sample was used, as a small concentration-dependent amount of dodecamer (< 3% at 10 mg/mL) was detectable from deconvolution of the higher concentrations.

Isotropic and anisotropic IPAP spectra for RDC data were collected in 50 mM sodium-phosphate pH 6.6, 2mM DTT at 298 K in the absence and presence of 20 mg/mL *Pf1* phage as the alignment medium. The lowest energy initial trimer structure (from above) was used as the starting point for simultaneous refinement against NOE, dihedral, RDC, and SAXS restraints in Xplor-NIH [78]. Agreement between measurements and values calculated from the structure was assessed using the DC server (https://spin.niddk.nih.gov/bax/nmrserver/dc/). Following refinement, the AT_3_ structure ensemble was validated using protein structure validation server (PSVS) [79] and Procheck-NMR [80]. Final refinement and validation statistics are provided in Table 1.

### NMR relaxation data

The ^15^N relaxation data were recorded on a Bruker DRX 600 MHz spectrometer at 328 K using water flip-back and 3-9-19 WATERGATE solvent suppression, sensitivity improvement, and gradient coherence selection [67]. All relaxation data were recorded with a spectral width of 8389.26 Hz sampled over 1024 complex points in ω_2_ (^1^H) and a spectral width of 1763.75 Hz sampled over 64 complex points in ω_1_ (^15^N), with 16 scans per transient except for the NOE which was collected with 64 scans per transient, and with a recovery delay between transients of 3 seconds. The recovery delays for *R*_1_ measurements were (100, 200, 400, 800, 1200, 1600, 2000, 2400 ms for AT_12_ and 10, 100, 250, 400, 550, 700, 850, 1250, and 2000 ms for AT_3_); relaxation delays for *R*_2_ measurements were (0, 9, 16.2, 23.4, 30.6, 37.8, 41.4, 45, 52.2, and 59.4 ms for AT_12_ and 0, 17, 33.9, 50.9, 67.8, 84.8, 101.8, 135.7, 169.6 ms for AT_3_). The NOE values were determined as the ratio of the peak intensities from two interleaved spectra recorded with and without a 3 second ^1^H presaturation period consisting of a series of 120° pulses separated by 5 ms delays. Relaxation rates were obtained by non-linear least-squares fitting measured peak heights with mono-exponential functions; uncertainties in peak heights were estimated from the standard deviation of the noise.

### Rotational diffusion

Overall rotational correlation times, τ_m_, and rotational diffusion tensors were determined experimentally from R_2_/R_1_ ratios [50] utilizing only R_2_/R_1_ ratios for residues with relaxation rates within one standard deviation of the 10% trimmed mean value. These calculations showed no statistical significance for fitting the NMR data to an axially symmetric or fully anisotropic diffusion tensor. Therefore, an isotropic diffusion tensor with a τ_m_ of 25 ns was used for the model-free calculations of the dodecamer. This was supported by calculations of the hydrodynamic parameters by HYDRONMR [81,82] based on the dodecameric structure 2BX9 that also predict nearly isotropic tumbling for AT_12_. The calculations based on the R_2_/R_1_ ratios of AT_3_, however, indicated a statistically significant improvement for the axially symmetric model over the isotropic model. This was similarly supported by HYDRONMR calculations that yield an anisotropic diffusion tensor with two similar and one unique axis. Therefore, an axially symmetric diffusion tensor from the R_2_/R_1_ ratios was used for the model-free calculations of AT_3_.

### Model-free analysis

Model-free analysis for both AT_3_ and AT_12_ [53] was performed using the program TENSOR v. 2 [83], which was recompiled to allow for input of up to 1000 residues and a CSA of -172 ppm [84]. Five motional models with up to three parameters describing the internal motions were used: (1) *S*^2^ (2) *S*^2^ and *t*_e_ (3) *S*^2^ and *R*_ex_ (4) *S*^2^, *t*_e_, and *R*_ex_ (5) *S*^2^, *S*^2^_f_, and *t*_e_. Standard errors in the model-free parameters were obtained from Monte Carlo simulations using 1000 samples. Model fitting was performed using TENSOR v. 2 via a modified step-up hypothesis testing method [54].

### Observation of the pH-dependent AT3-AT12 equilibrium via SAXS

The program CRYSOL [45] was used to generate predicted scattering profiles AT_3_ structure, the *B. subtilis* AT dodecamer retrieved from the RCSB (3BX9), and the *B. licheniformis* high-pH “inverted” dodecameric isoform (3LD0). Given the structural similarity of the AT trimer component between *B. subtilis* and *B. licheniformis* at nominal SAXS resolution (10-20 Å), the *B. licheniformis* dodecamer coordinates were used without modification. All three profiles were normalized to the square of the input structure’s molecular weight. The solvent density and contrast of the hydration shell were determined by fitting the scattering of the AT trimer structure against an empirical scattering profile of 5 mg/mL *f*AT in CRYSOL. The empirical scattering data from mature AT at 16 different pH and concentration conditions was fit with the program OLIGOMER [44], using the three predicted scattering profiles. An additional systematic term was also used to correct for baseline offset arising from incomplete buffer subtraction/mismatch.

### Analytical ultracentrifugation

AT stock solutions were split and dialyzed against sample buffers at pH 7, 7.25, 7.5, and 8, consisting of 50 mM sodium phosphate, 100 mM NaCl, 0.5 mM tryptophan, and 0.02% (w/v) NaN_3_. Sedimentation samples were subsequently made from the concentrated stock solutions with the corresponding dialysis buffer and allowed to equilibrate overnight. To account for the decreasing self-affinity at increasing pH, samples at pH 7-7.25 were made to 250, 125, and 65 µM AT monomer concentration, and at pH 7.5-8, to 500, 250, and 125 µM. Dilutions were performed on a gravimetric balance accurate to 0.2 mg, with an assumed solution density of 1.0 g/ml. Sedimentation velocity data were obtained at 25 °C using a Beckman Coulter ProteomeLab XL-I ultracentrifuge equipped with an 8-position rotor and double-sector cells with sapphire windows. Matched dialysis buffer was present in the reference sector. Data were recorded using interference optics with a scan interval of one minute and 200 scans per cell at a rotor speed of 50,000 rpm. The position of the meniscus and cell bottom were determined for each cell by fitting a sedimentation coefficient distribution and uniform frictional value f/f_0_ in SEDFIT [85] at a sampling interval of 10 to 20 points per Svedberg. Time and radial invariant noise were removed from each interference scan. The buffer density ρ and viscosity η under these conditions were calculated by SEDNTRP [86] to be 1.01260 g/mol and 1.0520 cP, respectively.

Determination of kinetic parameters for the AT_3_ ↔ AT_6_ ↔ AT_12_ model in SEDPHAT [43] was performed by globally fitting 200 interference scans from each set of AT concentrations at one pH (600 scans total). In addition to model kinetic parameters, the fringe shift coefficient, apparent sedimentation coefficients for AT_3_, AT_6_, and AT_12_, as well as the molar mass for the trimer were also optimized. pH 8 data were fit first, using the HYDROPRO [87]-predicted sedimentation coefficients of 2.1 S and 4.1 S at 25 °C for AT_3_ and AT_12_, respectively, as starting sedimentation coefficients. The AT_6_ starting sedimentation coefficient was set to 3 S. Initial parameters were 10^5^ M^-1^ for *k*_1_, 10^-3^ s^-1^ for *k*_-1_, 10^6^ M^-1^ for *k*_2_, and 10^-4^ s^-1^ for *k*_-2_, as these provided reasonable predicted interference curves. The apparent molar weight of AT in the experiment was also optimized, using the expected mass of the trimer as a starting value. A reasonable starting point for the molar absorptivity was determined by manual iterative evaluations of the model using different values until a fringe shift of appropriate magnitude was achieved. (Uncertainty analysis in the kinetic parameters was attempted through Monte-Carlo estimation of errors, but after 1000 evaluations no variance in the fitted parameters was detected.)

### Interfaces

Solvent accessible surface (SAS) area burial between protomers was computed using PyMOL 2.5.2 (Schrödinger, LLC) using a dot density of 3. Burial was computed from difference between the SAS of components and their complex, using PyMOL get_area. SAS burial for folding of the protomer was estimated from multiple conformations using PROSA [88]. Similar interface and surface areas were obtained using PISA [89]

## Acknowledgements

This work was supported by grants from the NIH to MPF and PG (GM077234, GM062750, GM120923). SAXS data were collected at the SIBYLS beamline at the Advanced Light Source (ALS), a national user facility operated by Lawrence Berkeley National Laboratory on behalf of the Department of Energy, Office of Basic Energy Sciences, through the Integrated Diffraction Analysis Technologies (IDAT) program, supported by DOE Office of Biological and Environmental Research. Additional support comes from the National Institute of Health project ALS-ENABLE (P30 GM124169) and a High-End Instrumentation Grant S10OD018483. Charles Cottrell (deceased) and Chunhua Yuan (CCIC) provided support for NMR experiments, Charles D. Schwieters for help with XPLOR-NIH and the members of the Foster laboratory for helpful discussions.

